# A novel technology for objective, accurate and non-invasive early diagnosis and monitoring of Alzheimer’s disease in clinics and clinical trials

**DOI:** 10.1101/790469

**Authors:** Kaveh Vejdani, Thomas Liebmann, Nicolas Pannetier, Elham Khosravi, Mohsen Yousefnezhad, Pavan Krishnamurthy, Nima Farzan, Cameron Bahar, Ahmad Salehi, Hesaam Esfandyarpour, Padideh Kamali-Zare

**Affiliations:** Darmiyan, Inc., 1 Sansome Street, Suite 3500, San Francisco, CA 94104, USA

**Author notes:** Corresponding author: Kaveh Vejdani, MD ‖ Chief Medical and Technology Officer, Darmiyan, Inc. ‖. Shared first authors (Kaveh Vejdani, Thomas Liebmann, Nicolas Pannetier).

**Keywords:** Dementia, Alzheimer’s disease, mild cognitive impairment, magnetic resonance imaging, MRI, early detection, diagnosis, prognosis, machine learning, ML, artificial intelligence, AI, clinical trials, patient selection, drug effect monitoring

## Abstract

We have developed a novel medical image processing technology for objective, accurate, and non-invasive diagnosis, prognosis, and monitoring of Alzheimer’s disease using standard clinical brain MRI (magnetic resonance imaging) and basic clinical cognitive assessment. The technology is a robust, highly accurate, fully automated, high-throughput, cloud-based platform, which operates in a fully integrated and controlled environment. In ***diagnosis mode***, our technology performs with 91% balanced accuracy on blind testing to diagnose the current neurocognitive status, and in ***prognosis mode*** *or early detection at mild cognitive impairment (MCI) stage* with 88% balanced accuracy on blind testing to predict progression from MCI to Alzheimer’s dementia (AD) within 5 years. Such prognostic capability is currently non-existent, even in specialty clinics and hospitals, a major factor in Alzheimer’s clinical trial failures. The algorithm’s diagnostic certainty precisely mirrors the diagnostic confidence of an expert cognitive neurologist for both MCI (Spearman’s Rho = 1) and AD (Rho = 1). In addition to widespread clinical applications, this novel technology can enable correct patient selection and therapeutic effect monitoring in clinical trials of Alzheimer’s disease, the crucial elements to finding a cure.

## Introduction

Dementia is the largest and most expensive healthcare challenge of our generation and likely the next. Fifty million people worldwide suffer from dementia, 60%-80% of which is due to Alzheimer’s disease. The global cost of dementia care has already reached a trillion dollars [1] and is projected to double by 2030 as the baby boomer generation is aging [2]. As families, governments and private insurance providers are heavily burdened by the increasing cost of dementia care, scientists and clinicians remain uncertain about the mechanisms underlying neurodegeneration and how to detect early, prevent, slow down, or stop it. Figure 1 depicts the course of Alzheimer’s disease with emphasis on the lack of an early detection method.

**Figure 1.**
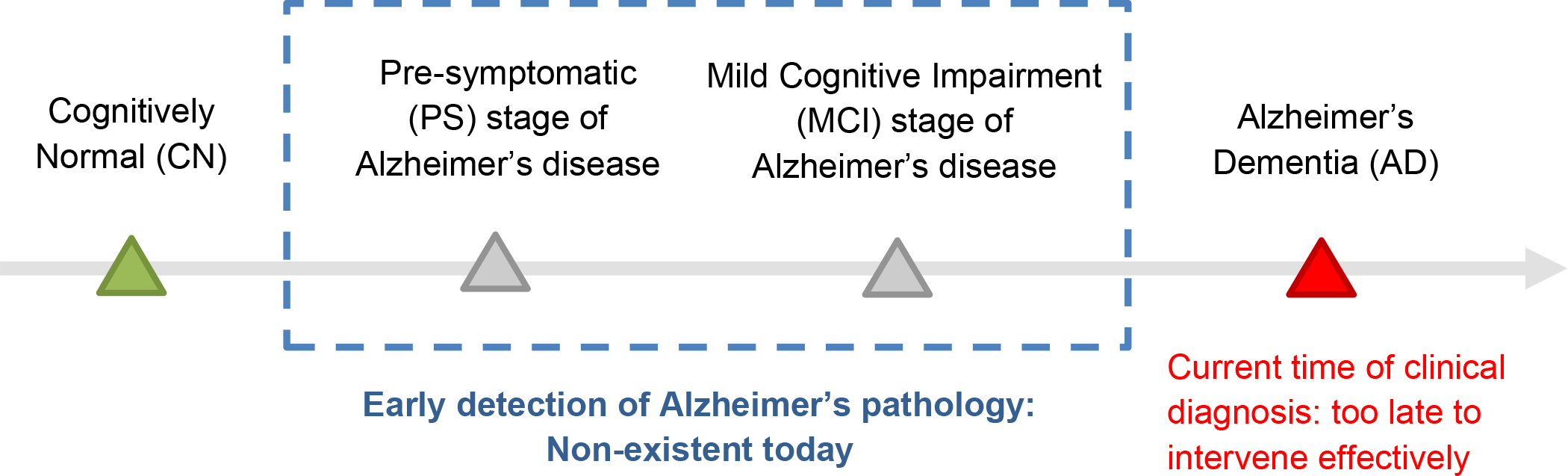
Stages of Alzheimer’s disease, the leading cause of dementia. Early detection of Alzheimer’s pathology in PS and MCI stages is missing in clinical practice and clinical trials today. Note that MCI may also be caused by brain diseases other than Alzheimer’s.

Initially promising drug candidates keep failing one after another in large clinical trials [3]. The reasons for these failures are shown in Figure 2 [4]. These include the lack of accurate and objective methods to 1) identify who should be selected for clinical trials, those who definitely have Alzheimer’s disease; 2) identify those at earlier stages of the disease who can potentially benefit from promising therapeutics; 3) stratify different sub-types of Alzheimer’s disease based on localization of pathology in the brain and to match each trial population with the drug with the most appropriate mechanism of action, and 4) monitor drug effect during clinical trials by measures that are agnostic to the cause of the disease and reflect the objective health of brain tissue in alignment with expected biological and clinical gain in cognition and disease modification.

**Figure 2.**
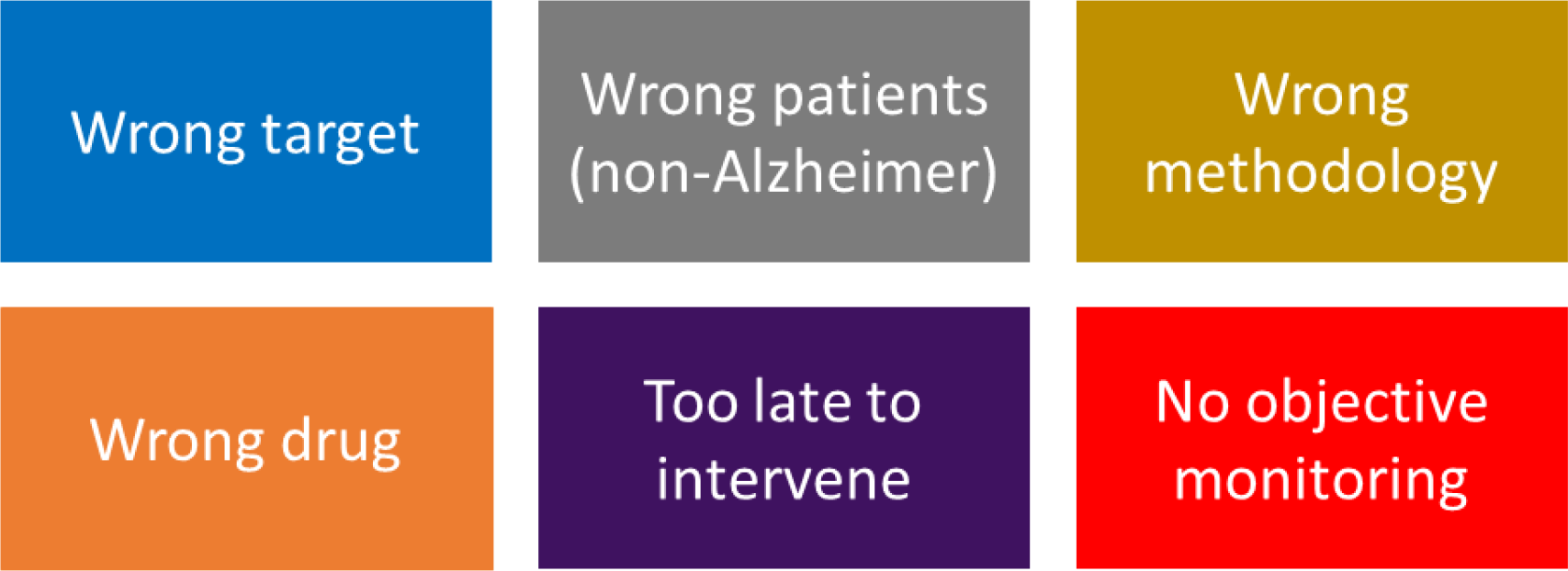
Reasons why Alzheimer’s drug trials keep failing [4].

Cognitive testing and medical history taking that are currently routinely used in clinics and hospitals reveal a lot of information about the current cognitive status of the brain and therefore are used as the basis for the clinician’s assessment. However, they remain inaccurate and unreliable about the prognosis or future cognitive status of the patient. Even for evaluation and diagnosis at the time of the patient’s visit, there is a great amount of subjectivity both in cognitive assessment and in the clinician’s judgement [5].

By the time there is a significant amyloid-beta plaque burden in the brain, amyloid-targeting PET (Positron Emission Tomography) tracers can reveal and quantify the plaque burden *in vivo* [6–8]. Amyloid-beta, although a classic hallmark of Alzheimer’s disease, is a non-specific biomarker which can accumulate in normal aging and in other neurodegenerative and non-neurodegenerative diseases as well. Whether or not detecting amyloid plaque burden by PET will affect the clinical outcome remains to be seen in future studies [9]. Amyloid-induced neurotoxicity in Alzheimer’s disease seems to be irreversible, even after the amyloid plaques are completely wiped out with anti-amyloid therapeutics, raising doubts on the anti-amyloid treatment approach. Amyloid PET is an expensive imaging modality which is not readily available in hospitals around the world. Repeated Amyloid PET scanning subjects the individual to cumulatively significant amounts of X-Ray radiation during the CT (computed tomography) portion of the scan, in addition to the radiation coming from the injected radioactive tracer, F18-labeled anti-Amyloid antibody.

Alzheimer’s disease is generally considered as an amyloid-induced tauopathy [10]. By the time there is significant neurofibrillary tangle burden in the brain, tau-targeting PET tracers can reveal and quantify the tau load *in vivo* [6,7]. Whether or not abnormal tau protein is the right biomarker to target and whether removing tau pathology would modify the course of Alzheimer’s disease remains to be seen in future clinical trials. Tau PET, like amyloid PET is an expensive imaging modality which is not readily available in hospitals around the world. Similar to amyloid PET imaging, repeated tau PET scanning exposes the individual to cumulatively significant amounts of X-Ray radiation during the CT portion of the scan, in addition to the radiation coming from the injected radioactive tracer, F18-labeled anti-Tau antibody.

Loss of gray matter mass in later stages of neurodegeneration can be revealed by FDG (Fluoro-Deoxy Glucose) PET scan, which reflects reduced glucose metabolism in the affected brain regions. The pattern of hypometabolism on FDG PET scan is used clinically for differential diagnosis of the etiology of dementia [6,7]. Given that glucose metabolism is not specific to any brain disease, whether or not the gray matter metabolic rate can be used as a biomarker to monitor anti-Alzheimer treatment effect has not yet been established. Although in common clinical practice for four decades, predominantly for oncologic applications, FDG PET remains an expensive imaging modality, with availability limited mostly to industrial or developed countries. Repeated FDG PET scanning subjects the individual to cumulatively significant amounts of X-Ray radiation during the CT portion of the scan, in addition to the radiation coming from the injected radioactive tracer, F18-labeled glucose.

Amyloid and tau protein measurements in the cerebrospinal fluid (CSF) can be used as surrogates for amyloid and tau presence in the brain. Obtaining a CSF sample requires an invasive procedure called a lumbar puncture, which is costly and very inconvenient for the patient. Repetitive CSF sampling is impractical in both clinical and research settings.

Unlike PET imaging and CSF sampling, MRI has the advantage of being safe (no radiation or contrast agent injection), non-invasive, information-rich, and relatively inexpensive, at about 1/4 of the cost of PET. MRI can reveal detailed information about the brain tissue at millimeter scale, as is routinely evaluated by the clinical radiologist. It is the standard modality in clinical neuroimaging and MRI scanners are widely available throughout the world [11].

The development of machine learning (ML) and artificial intelligence (AI) methods for computer-aided diagnosis and prognosis of Alzheimer’s disease has been very active in the past decade [8,12]. This has been made possible by the joint effort of the scientific community and funding agencies to: 1) collect, curate and publicly release longitudinal databases focused on Alzheimer’s disease e.g. ADNI, OASIS, and AIBL, which include multiple data collection methods and modalities including cognitive tests, genetics, MRI, and PET, and 2) democratize machine learning techniques through standardization of training and evaluation and facilitated access to cutting-edge algorithms.

The earliest results of studying Alzheimer’s disease using neuroimaging data started to emerge in 2005 and the accuracy and robustness of the techniques kept increasing as larger datasets were made available and more advanced techniques were developed and employed. In the clinically useful and relevant context of predicting progression from MCI to AD (prognosis or early detection at MCI stage), performance accuracy has been reported to range between 65% to 80% with serious questions raised around the generalizability and applicability of such methods in the actual clinical setting [8]. Combining different modalities such as MRI, PET, and CSF biomarkers can be inconvenient and costly for patients as they have to go through several tests in order to get a reliable diagnosis and prognosis. Another important issue around ML/ AI algorithms is that they are mostly trained on data that is collected in a well-controlled research setting. This limits the performance accuracy and application of such trained algorithms on blind samples coming from a variety of external sources, representing the real-world situation.

## BrainSee, Darmiyan’s novel technology

Our team at Darmiyan, Inc. has developed a novel, ML-powered, proprietary technology, BrainSee, [13] that has incorporated the following modules in a fully automated, integrated, and controlled environment:

1. **Data curation and standardization module:** Pre-processes raw brain MRI (DICOM format) and prepares it for subsequent processing steps
2. **Neuroscience module:** Maps MRI features to key biological and physiological features of human brain tissue in every voxel
3. **Clinical data module:** Curates the additional demographic and clinical metadata available for each subject at each data point
4. **Scoring module:** Combines the output of module 2 in various brain regions in a unique way and generates a single score reflective of the health and disease status of the brain
5. **ML/ AI module**: Uses ML/ AI to empower diagnosis and prognosis capabilities

The current version of the technology, BrainSee v1.0, is a cloud-based platform that inputs standard clinical brain MRI and basic clinical cognitive assessment and outputs 1) Diagnostic decision: CN, MCI, or AD, with a separate probability score for each class, and 2) Prognostic decision: Stable MCI or Converter MCI, with probability of conversion to AD within the next 5 years. BrainSee does not require any specialized scan parameters, contrast agent, or additional hardware.

## Methods for validating the performance of BrainSee v1.0

### Diagnosis mode

#### Data source

23 clinical sites from ADNI, 3179 data points

#### Input

Age, Sex, Education (years), MMSE (total score), CDR (all boxes and global)

#### Output

Probability of CN, probability of MCI, probability of AD, diagnostic decision (label corresponding to the highest probability). Probability of each class also represents the certainty or confidence in that diagnostic class decision.

#### Ground truth for evaluation

The diagnostic label provided by the clinician (either CN, MCI, or AD). The clinician’s diagnostic decision was based on the patient’s history, comorbidities, medications, and cognitive and functional assessments such as MMSE, CDR, FAQ, and ADAS. The clinician generally has access to more detailed information than what is provided to BrainSee.

#### Clinician’s diagnostic confidence

Each clinician scored their diagnostic confidence for each time point on a scale of 1 to 3, where 1 = uncertain or low confidence, 2 = moderate confidence, and 3 = high confidence. Clinician’s diagnostic confidence information was available only in a subset of the data, coming from ADNI1 phase.

#### Analysis

BrainSee was fully blind to the data source. No data point was previously used in the development or internal testing of BrainSee. To evaluate the diagnostic accuracy, BrainSee’s labels were matched against the clinician’s labels. Performance (recall) was reported separately for each class (CN, MCI, AD). Balanced accuracy was calculated as the arithmetic mean of all class recalls. To evaluate diagnostic certainty, BrainSee’s certainty was correlated with the clinician’s confidence using Spearman’s rank-order correlation coefficient (Rho).

### Prognosis mode

#### Data source

20 clinical sites from ADNI, 411 data points

#### Input

Age, Sex, Education (years), MMSE (total score), CDR (all boxes and global), MRI scan (T1, T2, and diffusion-weighted sequences)

#### Requirement

All subjects had amnestic MCI at the time of MRI scans

#### Output

Probability of progression from MCI to AD within 5 years of the MRI scan; Prognostic decision: Stable MCI if probability of progression is < 50%, or Converter MCI if probability of progression is > 50%.

#### Ground truth for evaluation of progression from MCI to AD

The future clinical assessment at the 5-year follow-up as decided by an ADNI clinician was set as reference for ground truth. Such decision is based on cognitive and functional assessments at the time of the future visit, i.e. up to 5 years after the algorithm makes its prediction. MCI stability is defined as non-conversion to AD at least 5 years after the scan (minimum 5 years of staying MCI), and progression to AD was ascertained by conversion within 5 years of the MRI scan.

#### Analysis

BrainSee was fully blind to the data source. No data point was previously used in the development or internal testing of BrainSee. To evaluate the prognostic accuracy, BrainSee’s prediction of future cognitive status was compared against the clinician’s diagnostic label at a clinical visit 5 years after the MRI scan.

## Results of BrainSee v1.0 performance

### Diagnosis mode

BrainSee’s diagnosis label matches that of a qualified clinician with 91% balanced accuracy. BrainSee’s diagnostic confidence matches that of a qualified clinician for MCI and AD classes: MCI class Spearman’s Rho = 1 (sample size n=529), AD class Rho = 1 (n=435). Performance accuracy (recall) results on 3179 blind data points:

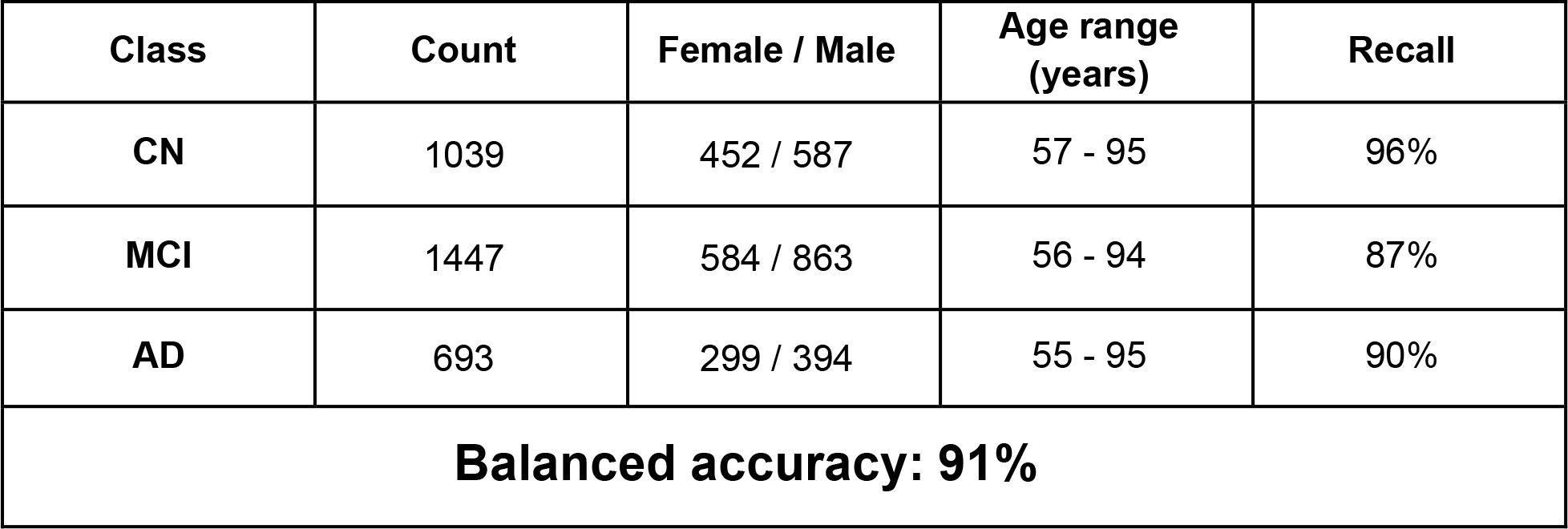

| Class | Count | Female / Male | Age range (years) | Recall |
| --- | --- | --- | --- | --- |
| CN | 1039 | 452 / 587 | 57 - 95 | 96% |
| MCI | 1447 | 584 / 863 | 56 - 94 | 87% |
| AD | 693 | 299 / 394 | 55 - 95 | 90% |
| **Balanced accuracy: 91%** |  |  |  |  |

### Prognosis mode

BrainSee predicted clinical progression from MCI to dementia within 5 years of the MRI scan with 88% balanced accuracy. Performance accuracy (recall) results on 411 blind data points:

| Class | Count | Female / Male | Age range (years) | Recall |
| --- | --- | --- | --- | --- |
| Stable MCI | 139 | 37 / 102 | 56 - 89 | 85% |
| Converter MCI | 272 | 122 / 150 | 57 - 91 | 91% |
| **Balanced accuracy: 88%** |  |  |  |  |

Given the very strong correlation between a clinician’s diagnostic confidence and BrainSee’s diagnostic certainty, BrainSee’s certainty can be used as a surrogate to the clinician’s confidence. Removing clinically uncertain cases, i.e. cases where the presence of dementia at 5-year follow-up is uncertain, results in up to 5% gain in performance accuracy.

### Sample report

An example output report of BrainSee, from a 79-year-old female MCI patient, is shown below.

BrainSee takes the patient’s age, sex, cognitive assessment, and MRI scan files as input, and generates the cognitive diagnosis and prognosis as output. The prognostic score, i.e. the chance of progression from MCI to AD, is shown as a red dot overlaid on a scatter plot background of all other previously tested subjects. Orange dots represent converter cases, i.e. those who progressed from MCI to AD in 5 years, and blue dots represent stable (non-converter) cases, i.e. those who stayed MCI for 5 years or longer. Gray dots represent misclassified cases.

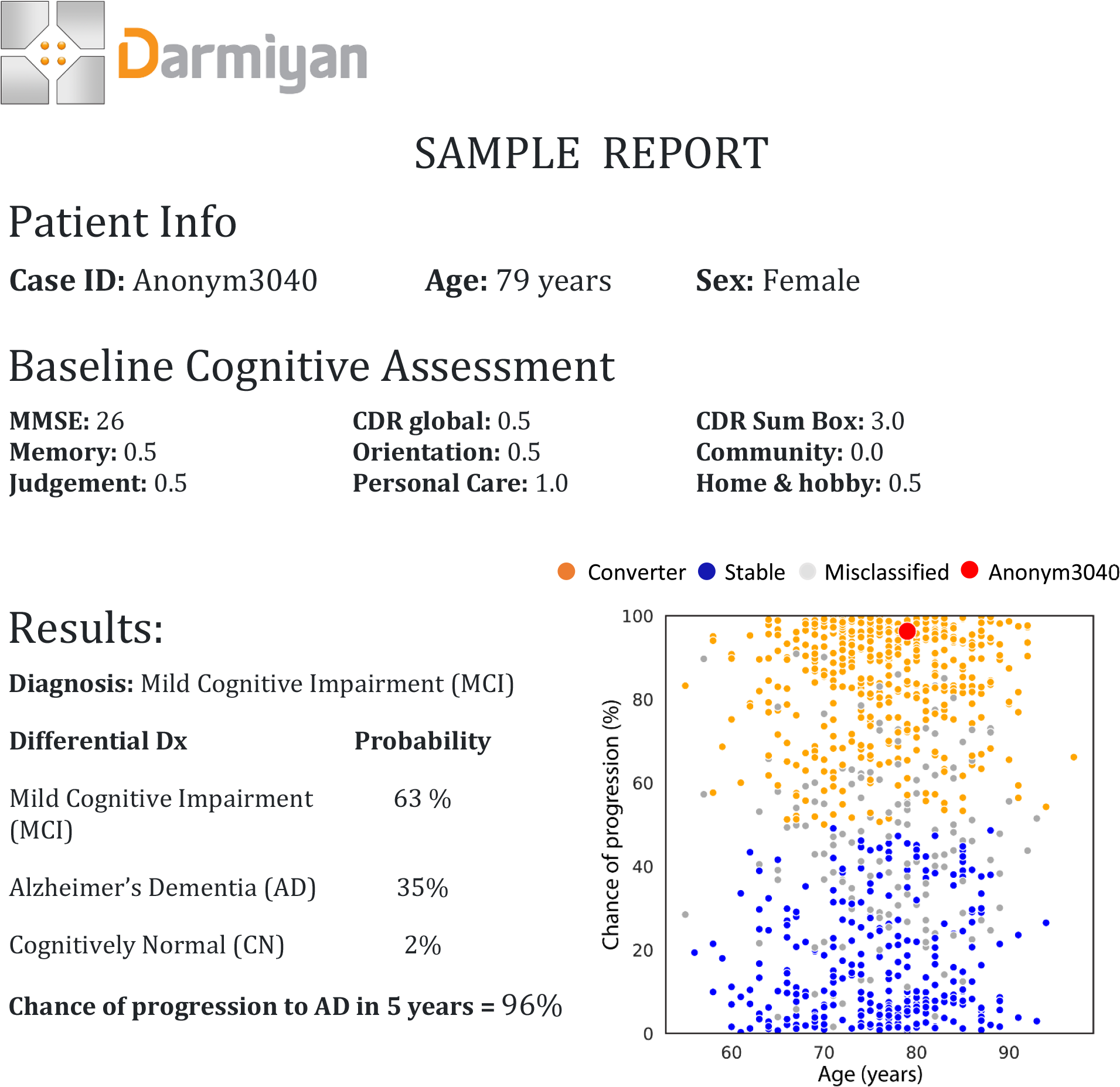

## Discussion & conclusion

Early diagnosis of Alzheimer’s disease, even without a treatment, leads to significant cost savings of $7T in the current US population (Figure 3) mainly through better managed comorbidities and reduced hospitalizations [1]. The major roadblock in finding a treatment for Alzheimer’s disease is the lack of an accurate, reliable, and objective method for early detection of the disease and for monitoring therapeutic efficacy. Currently, there is no cure for Alzheimer’s disease even when detected. By the time people are clinically diagnosed today, it is too late to intervene effectively as the damage to the brain tissue has progressed too far to be reversed. All disease-modifying clinical trials have failed to date. Each trial costs the pharmaceutical industry on average $5.7 Billion for drug development and testing [14].

**Figure 3.**
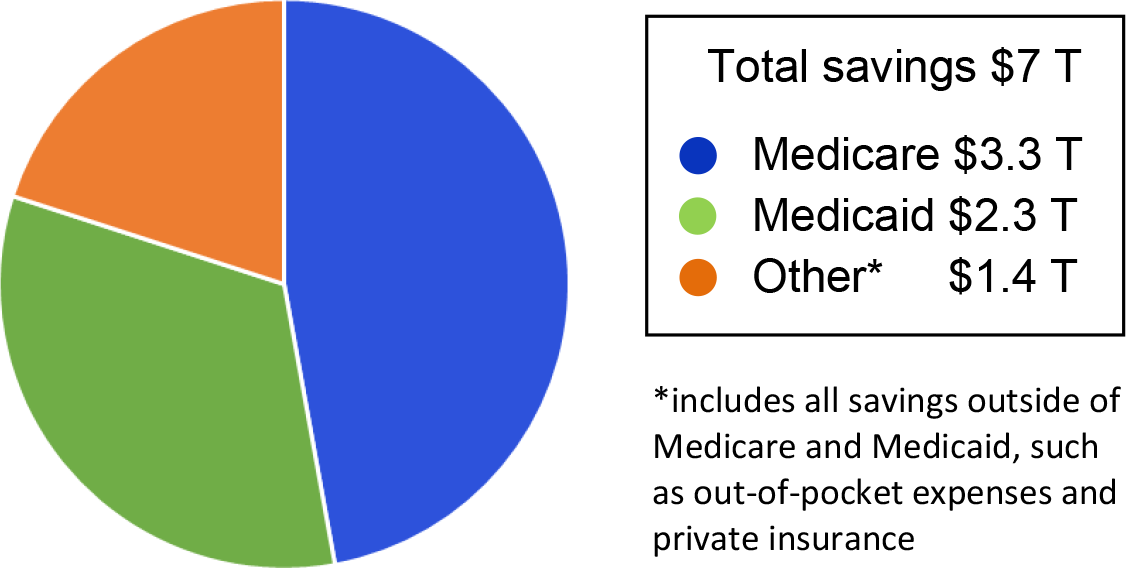
Cost savings associated with early diagnosis of Alzheimer’s disease. More than half of the cost savings was attributed to lower inpatient hospital costs, modified from [1].

A doctor’s diagnosis of Alzheimer’s disease determines the course of a patient’s clinical management and potential decision for enrollment in a clinical trial. Patients and their families discuss care and life planning based on the diagnostic information provided to them by the clinician. When the clinician is uncertain, patients and their families remain uncertain and worried [1,2,15]. Even the most widely accepted guidelines in the medical community leave the clinician with great uncertainty in their subjective assessment of MCI and AD [16].

Clinical uncertainty is not limited to differentiating MCI from dementia diagnosis [15–17]. Even with a fairly certain clinical diagnosis of MCI, predicting the future course (prognosis) of the underlying abnormality in terms of progression, stability, or reversal, remains uncertain [15]. In medical practice, such prognostic uncertainty creates additional worries and anxieties in the patient and their families. It also makes it difficult for the clinician to decide confidently on an appropriate treatment. In clinical trials, prognostic uncertainty makes it extremely difficult to recruit the appropriate patients to power the study properly and demonstrate statistically significant difference between the treatment arm and placebo arm.

Our novel solution, BrainSee, provides objective, accurate and non-invasive early diagnosis and monitoring of Alzheimer’s disease for clinics and clinical trials. It removes subjectivity and uncertainty from clinical assessments of MCI and AD and enables correct patient selection in clinical trials of Alzheimer’s disease (Figure 4).

**Figure 4.**
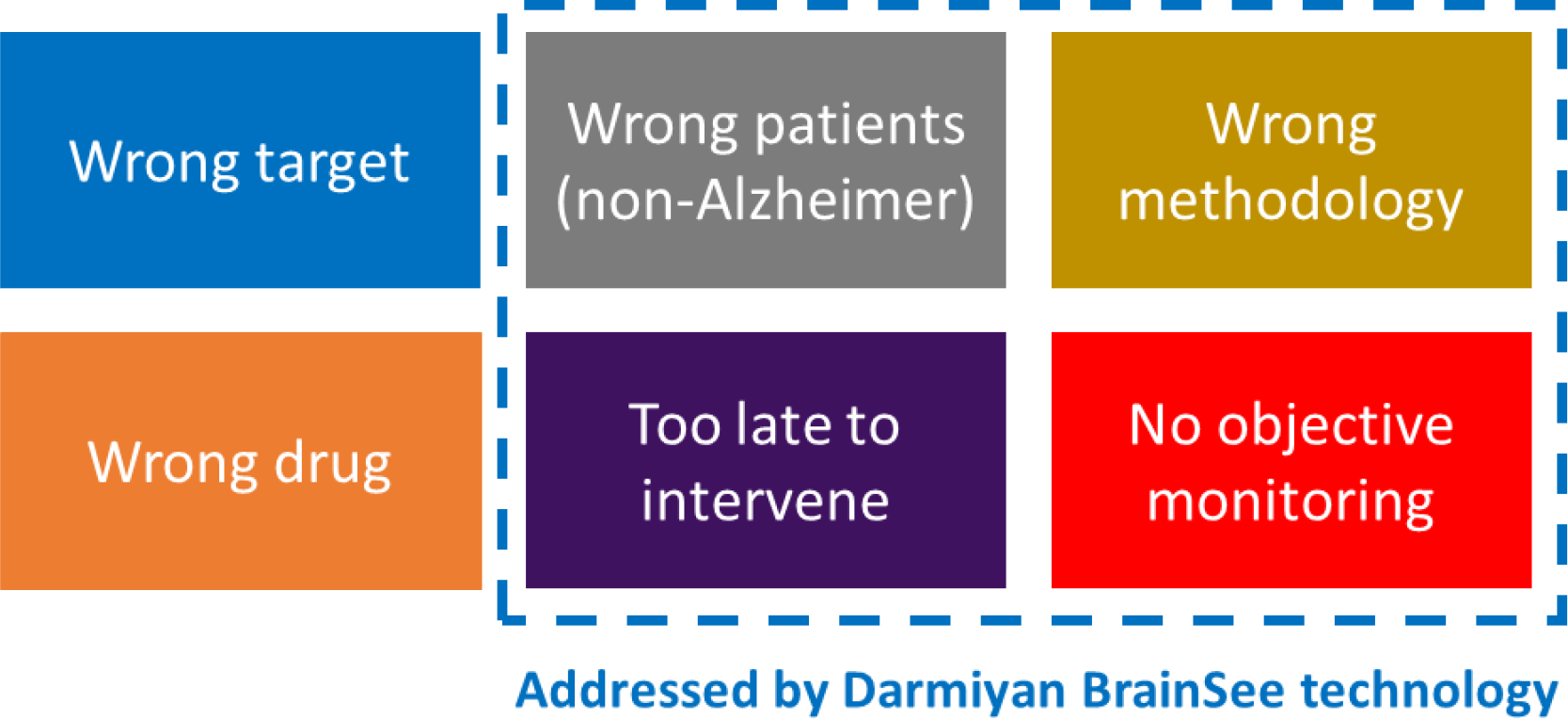
Four of the major reasons of Alzheimer’s disease clinical trial failures are addressed by Darmiyan’s novel BrainSee technology.

BrainSee is a fully automated, cloud-based platform, which operates in a fully integrated and validated environment, a critical feature lacking in most other medical image processing technologies. It performs with high accuracy (~90%) and is designed for robust, high-throughput analysis of images from different sources. It operates in diagnosis and prognosis modes:

In **diagnosis mode**, BrainSee provides an accurate diagnosis of the cognitive state at the time of clinical evaluation. It takes age, sex, years of education and standard cognitive test scores (MMSE & CDR) as input data and generates a probability score for each cognitive state (CN, MCI, and AD), as well as a diagnostic decision. These output scores have the following applications:

1. For clinical use: As novel biomarkers for monitoring the brain health status, MCI severity and AD diagnosis certainty
2. For clinical trials: AD probability score can be used as a novel “disease modification” biomarker for therapeutic effect monitoring
3. For research use: To provide diagnostic labeling consistency and reliability across different sites and multiple data sources
In **prognosis mode**, BrainSee provides accurate detection of Alzheimer’s disease years before confirmation is possible in clinics. It takes the same input data as the diagnosis mode, plus the addition of standard brain MRI, which is already a standard, well-established imaging modality and integrated in the clinical workflow around the world. BrainSee prognosis generates the probability score for progression to dementia versus stable cognitive impairment. This output score has the following applications:

1. For clinical use: Early detection of Alzheimer’s disease at the MCI stage
2. For clinical trials: A novel “disease modification” biomarker for therapeutic effect monitoring
3. For research use: Standard framework to test various other research hypotheses

With these unique, novel features, BrainSee can solve the limitations in clinical practice and clinical trials of Alzheimer’s disease. Our upcoming paper will focus on the results of a simulated clinical trial enrichment and monitoring capability using BrainSee technology.

## Darmiyan, The Company

Darmiyan Inc. is a biotechnology company based in San Francisco, California, that was incorporated in September 2016. A team of neuroscientists, clinicians, data scientists, physicists, biophysicists, and mathematicians work together in Darmiyan to address the global need for objective and accurate early diagnosis of Alzheimer’s disease and to enable finding a treatment. The Darmiyan team have over four decades of combined neuroscience research and clinical experience, gained in major academic centers and industry laboratories and hospitals. We are driven to meet the challenge of tackling one of the last remaining frontiers in biomedical research, the human brain. By offering new non-invasive methodologies to study the brain in health and disease, we seek to bring clarity and advancement to the field.

## Data acknowledgement

Data used for blind testing in this article were obtained from the Alzheimer’s Disease Neuroimaging Initiative (ADNI) database (adni.loni.usc.edu). As such, the investigators within the ADNI contributed to the design and implementation of ADNI and/or provided data but did not participate in analysis or writing of this report. A complete listing of ADNI investigators can be found at: http://adni.loni.usc.edu/wp-content/uploads/how_to_apply/ADNI_Acknowledgement_List.pdf

## Abbreviations

CN: Cognitively Normal
MCI: Mild Cognitive Impairment
AD: **Alzheimer’s Dementia**
MMSE: Mini Mental State Exam
CDR: Clinical Dementia Rating
FAQ: Functional Assessment Questionnaire
ADAS: Alzheimer’s Disease Assessment Scale
MRI: Magnetic Resonance Imaging
PET: Positron Emission Tomography
CT: Computed Tomography
CSF: Cerebro-Spinal Fluid
ML: Machine Learning
AI: Artificial Intelligence
ADNI: Alzheimer’s Disease Neuroimaging Initiative
OASIS: Open Access Series of Imaging Studies
AIBL: Australian Imaging, Biomarkers and Lifestyle study

## Note

Throughout this article, the term “Alzheimer’s disease” is used as a general term to refer to all stages of the disease, including the asymptomatic stage, mild cognitive impairment stage, and dementia stage. The AD abbreviation specifically refers to the dementia stage of Alzheimer’s disease.

## Acknowledgements

We would like to thank all our investors, clinical partners and advocates who have supported the Darmiyan team and made our progress possible. Special thanks go to the Alzheimer’s Association, Canadian Center for Aging and Brain Health Innovation (CABHI), Baycrest Institute, University Health Network (UHN), Hamilton Health Sciences (HHS), Huntington Medical Research Institute (HMRI), Banner Alzheimer’s Institute, FlyWheel Exchange, Enzyme, IT-Farm, Dr. Enke Bashllari, Arkitekt Ventures, Soma Capital, Mr. Hamid Moghaddam, Mr. Soroush Kaboli, Mary Furlong & Associates, MedTech Innovator, SkyDeck UC Berkeley, Y Combinator, and TEDMED.

In memory of the late Professor Paul Greengard (Nobel Laureate in Physiology/ Medicine 2000), our Scientific Advisory Board member.

Data used for blind testing in this article were obtained from the Alzheimer’s Disease Neuroimaging Initiative (ADNI) database (adni.loni.usc.edu).

## Author contributions

BrainSee technology design and development: KV, TL, NP, EK, MY, HE, PKZ

Image analysis, machine learning pipelines and automation: NP, TL, EK

Data analysis: TL, NP, EK, KV

Manuscript: KV, PK, EK, PKZ

Technical and clinical feedback: HE, AS, PK, CB, NF

Directing the project: PKZ

## Author Information

All authors are employees, directors, advisory board members, or contractors of Darmiyan, Inc.

